# PHYLOSCANNER: Inferring Transmission from Within‐ and Between-Host Pathogen Genetic Diversity

**DOI:** 10.1101/157768

**Authors:** Chris Wymant, Matthew Hall, Oliver Ratmann, David Bonsall, Tanya Golubchik, Mariateresa de Cesare, Astrid Gall, Marion Cornelissen, Christophe Fraser

## Abstract

A central feature of pathogen genomics is that different infectious particles (virions, bacterial cells, etc.) within an infected individual may be genetically distinct, with patterns of relatedness amongst infectious particles being the result of both within-host evolution and transmission from one host to the next. Here we present a new software tool, phyloscanner, which analyses pathogen diversity from multiple infected hosts. phyloscanner provides unprecedented resolution into the transmission process, allowing inference of the direction of transmission from sequence data alone. Multiply infected individuals are also identified, as they harbour subpopulations of infectious particles that are not connected by within-host evolution, except where recombinant types emerge. Low-level contamination is flagged and removed. We illustrate phyloscanner on both viral and bacterial pathogens, namely HIV-1 sequenced on Illumina and Roche 454 platforms, HCV sequenced with the Oxford Nanopore MinION platform, and *Streptococcus pneumoniae* with sequences from multiple colonies per individual. phyloscanner is available from <u>https://github.com/BDI-pathogens/phyloscanner</u>.

## Introduction

The infectious transmission process imposes a hierarchical structure of relatedness on pathogen genomes. The genotype of an individual infectious particle is the result of both within-host evolution and transmission between hosts; a population sample collected from multiple hosts, with multiple genotypes for each host, therefore simultaneously encodes the history of both processes. Despite the existence of many tools for analysing pathogen genomes, none, to our knowledge, are specifically adapted to exploiting this hierarchical genealogical structure.

A central aim of infectious disease epidemiology is the identification of risk factors for transmission. The development of methods that use pathogen genomes to infer transmission events, along with their direction, is therefore a priority. A critical recent insight is that including multiple pathogen genomes per infected individual in such methods makes this inference easier: it is equivalent to the simpler process of inferring ancestry (Romero-Severson et al. 2016). Specifically, if a pathogen has passed from individual X to individual Y (either directly, or indirectly via unsampled intermediate individuals) then all the pathogen particles sampled from individual Y must be descended from the population of pathogen particles from individual X. Inferring ancestral states is a standard problem in population genetics for which many methods exist; the novel insight is that this standard approach may be used to infer the direction of transmission. We illustrate this in Figure 1.

**Figure 1:**
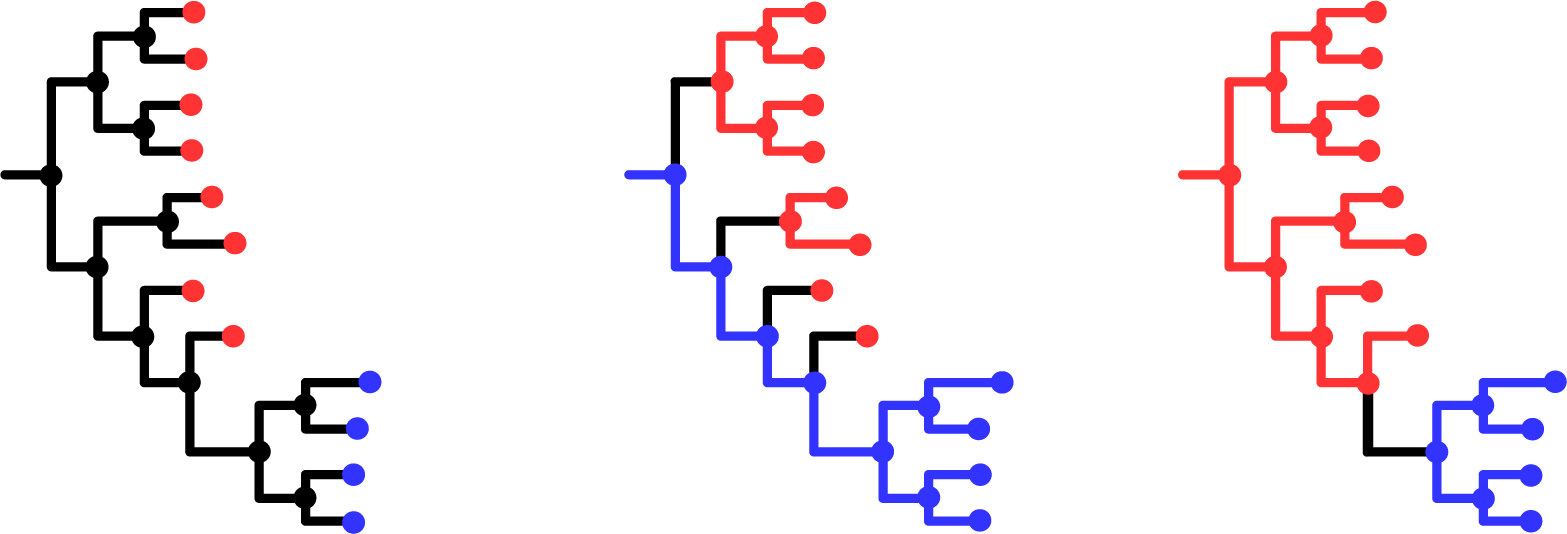
[pathogen transmission direction via ancestral state reconstruction. In the left-hand phylogeny, tips are labelled red or blue according to their state: in our case the state of interest is ‘in which individual was this pathogen found?’. This state is known for the tips, but can only be inferred for the internal nodes of the phylogeny: these represent coalescence events, ancestors of the pathogens we have sampled. A change in state corresponds to a change in the pathogen's host, i.e. to transmission, be it direct or indirect. The central phylogeny shows one possible ancestral state reconstruction for which the root of the tree is blue, meaning blue is ancestral to red. This requires at least four changes of state (shown with black branches) – four sampled lineages transmitted from blue to red. The right-hand phylogeny shows one possible ancestral state reconstruction for which the root of the tree is red, meaning red is ancestral to blue. This requires only one change of state – one sampled lineage transmitted from red to blue. Based on parsimony we would prefer the right-hand scenario.]

A frequently used approach in molecular epidemiology is to describe patterns of genetic clustering - who is close to whom. However, identifying transmission pairs or clusters without the ability to infer transmission direction - who infected whom - limits our ability to distinguish risk factors for transmission from those for simply acquiring the pathogen. One approach for inferring direction is to augment the sequence data with epidemiological data, and to couple phylogenetic inference with mathematical models of transmission, for example references (Volz and Frost 2013; Jombart et al. 2014; Hall et al. 2015; Didelot et al. 2017). However, this requires strong assumptions from the model. In addition epidemiological data, such as dates and location of sampling and reported contacts, are not always available, are subject to their own set of uncertainties and errors, or are sometimes regarded as too sensitive to link to pathogen genetic data.

Using multiple genotypes per host, and exploiting the link between transmission and ancestral reconstruction, therefore promises an alternative and potentially powerful approach to molecular epidemiology. Whilst several studies have used this idea to great effect on an ad hoc basis (Numminen et al. 2014; Worby et al. 2016), no systematic or automatic tool has been developed for this task.

Once multiple genotypes per host are included in a study, other questions present themselves naturally, for example identifying multiply infected individuals. These may be defined as individuals harbouring pathogen subpopulations resulting from distinct founder pathogen particles. Multiple infections may be clinically relevant, for example in the case of Human Immunodeficiency Virus 1 (HIV-1), dual infection is associated with accelerated disease progression (Cornelissen et al. 2012). Multiple infections also represent unique opportunities for pathogen evolution, especially for pathogens that recombine. Recombination between divergent strains accelerates the generation of novel genotypes, and so potentially novel phenotypes. The distinct pathogen strains in a multiple infection could have been transmitted simultaneously from the same individual (if that individual harboured sufficient within-host diversity), or sequentially – 'super-infection’ – with each strain perhaps originating from a different transmitter. For HIV-1, mathematical modelling has suggested that recombinants can reach high prevalence even when the possibility of super-infection is restricted to a short window after initial infection, and even when recombinants have no fitness advantage, if the epidemic is fuelled by a high-risk core group (Gross et al. 2004).

Molecular epidemiology is being transformed by the advent of next-generation sequencing (NGS; also called *high-throughput*) technologies (Goodwin et al. 2016). For many sequencing protocols applied to pathogens with extensive within-host diversity, such as HIV-1 and Hepatitis C Virus (HCV), the NGS output from a single sample can capture extensive within-host diversity. Zanini et al. (Zanini et al. 2015) inferred phylogenies from NGS *reads* - fragments of DNA - in windows along the genome for longitudinally sampled individuals infected with HIV-1, to quantify patterns of within-host evolution over time. Here our focus will be on cross-sectional datasets: by constructing phylogenies from NGS reads from multiple infected individuals at once, within-host and between-host evolution can be resolved.

We present phyloscanner: a set of methods implemented as a software package, with two central aims. The first is efficient computation of phylogenies with multiple genotypes per infected host, and the second is analysis of such phylogenies and inference of biologically and epidemiologically relevant properties from a set of related phylogenies. Multiple related phylogenies arise naturally, either by sampling different portions of a genome, or in representing uncertainty in phylogenetic inference (though bootstrapping, or sampling phylogenies from a posterior distribution, for example). phyloscanner automatically performs the following steps:

1. Inference of between and within-host phylogenies from NGS data in multiple windows along the pathogen genome (optionally skipped, if the user has such phylogenies already);
2. Identification and removal of likely contaminant sequences;
3. Quantification of within-host diversity;
4. Identification of multiple infections;
5. Identification of crossover recombination breakpoints in NGS genotypes;
6. Ancestral host-state reconstruction from multiple phylogenies;
7. Identification of transmission events from ancestral host-state reconstructions.

phyloscanner was intended for analysis of two distinct types of sequence data. Firstly for deep sequencing data, in which NGS has produced reads from the population of diverse pathogens represented in the sample. Secondly, for single-genome amplification (SGA), clonal sequencing or bacterial colony picks, whereby laboratory methods are employed to separate the genomes of individual pathogen particles prior to amplification and sequencing. Sequencing with primer IDs (Jabara et al. 2011) may in some cases produce similar results at reduced costs. We also considered haplotype reconstruction (Zagordi et al. 2011; Prabhakaran et al. 2014; Töpfer et al. 2014), i.e. bioinformatically inferring different haplotypes represented in the short reads of a mixed sample, but in our hands this approach did not yield satisfactory results (analysis not shown).

With SGA-style data, within- and between-host phylogenies can be directly inferred using standard methods, and therefore phyloscanner is not necessary for step 1 in the process described above. With deep sequencing data, reads for each sample must first be *mapped* (placed at the correct location in the genome); thereafter phyloscanner begins by aligning reads in windows of the genome that are matched across infected individuals, and inferring a phylogeny for each window (Figure 2).

**Figure 2:**
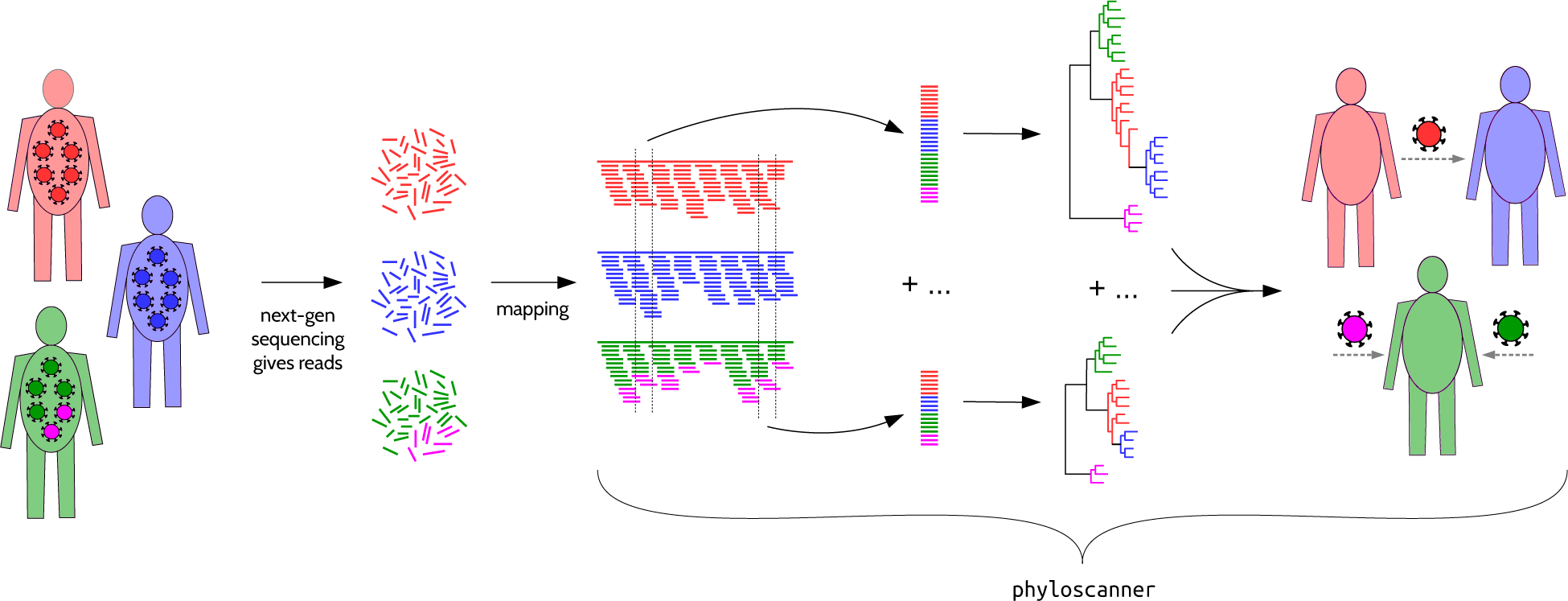
[phyloscanner schematic for whole-genome deep sequence data. In this schematic, pathogens are sampled from the population infecting three hosts. NGS deep sequencing produces reads, which are fragments of the genome sequence of one pathogen particle (after amplification if necessary). Mapping to a reference means aligning each read to the appropriate location in the genome; this must be done beforehand, as mapped reads are the inputs to phyloscanner. phyloscanner produces alignments of reads in sliding windows along the genome, automatically adjusting for the fact that the reference may be different for each sample. Phylogenies are inferred for each alignment. These phylogenies are analysed separately using ancestral host-state reconstruction (i.e. assigning hosts to internal nodes), and their information is combined to give biologically and epidemiologically meaningful summaries. For example here, we infer that the red individual infected the blue individual directly or indirectly, and the green individual has two distinct pathogen strains.]

## Results

The best way to illustrate phyloscanner is through examples. We chose five datasets illustrating different uses, pathogens, and sequencing platforms. We describe four in the main text, and one in the Supplementary Information. These are far from systematic samples or population surveys; they are small selections of infected individuals chosen to illustrate the different conclusions that can be drawn using phyloscanner. We leave the application of phyloscanner to large systematic population samples to future work.

Before presenting phylogenies for these data we introduce the term *host subgraph*. Host subgraphs result from ancestral host-state reconstruction: they are defined as connected regions of the phylogeny (tips and internal nodes, with the branches joining them) that have all been assigned the same host state (i.e., the host that pathogen was in). See supplementary section SI 1 for an explanation of the ancestral state reconstruction algorithm. Each subgraph can be shown with a solid block of colour corresponding to that host, uninterrupted by colouring associated with any other host. Figure 3 shows an example.

**Figure 3:**
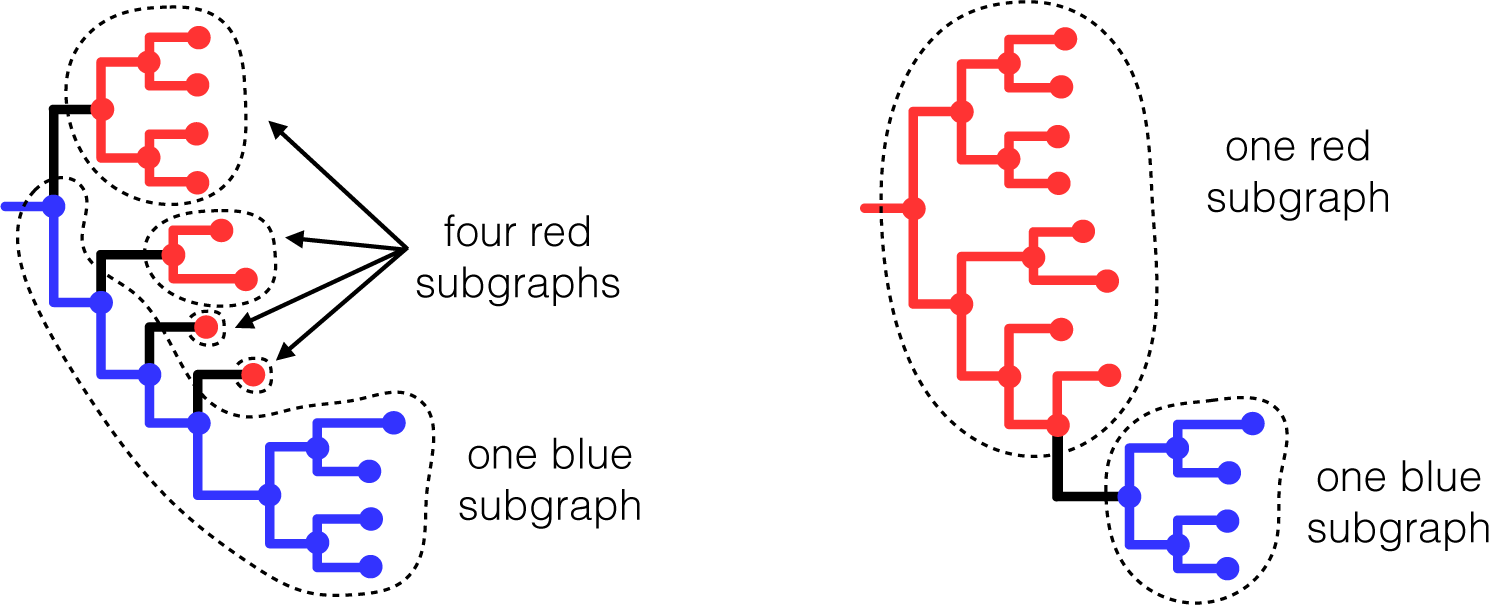
[subgraphs defined by a given ancestral state reconstruction. Here we show again the two different ancestral state reconstructions on the same phylogeny from Figure 1, this time illustrating the *host subgraphs* that these reconstructions define: connected regions of the phylogeny that have all been assigned the same state (blue host or red host). Note that the set of tips in a subgraph may or may not form a clade. In both of the above reconstructions, the blue tips are contained in one subgraph and form a monophyletic group (one clade), whereas the red tips form a polyphyletic group. The minimum number of clades needed to encompass all and only the red tips is four, coinciding with the four red subgraphs in the left-hand reconstruction.]

### Six illustrative HIV-1 infections, sequenced with Illumina MiSeq

We used phyloscanner to analyse data from the BEEHIVE project (*Bridging the Evolution and Epidemiology of HIV in Europe*), in which whole-genome samples from individuals with well-characterised dates of HIV-1 infection are being sequenced, primarily to investigate the viral-molecular basis of virulence (Fraser et al. 2014). We chose two groups of patients for detailed investigation (presented in this subsection and the next), that together demonstrate interesting features revealed by phyloscanner.

For the BEEHIVE samples, viral RNA was extracted manually from blood samples following the procedure of Cornelissen *et al.* (Cornelissen et al. 2016). The RNA was reverse transcribed and amplified using universal HIV-1 primers that define four overlapping amplicons spanning the whole genome, then sequenced using the Illumina MiSeq platform, following the procedure of Gall *et al.* (Gall et al. 2012; Gall et al. 2014). The resulting reads were mapped to a reference constructed for each sample using IVA (Hunt et al. 2015) and shiver (Wymant et al. 2016), producing input analogous to the illustration in Figure 2. See Materials and Methods for more detail.

These mapped reads were analysed with phyloscanner using 54 overlapping windows, each 320 base pairs (bp) wide, covering the whole HIV-1 genome (approximately 9200 bp long; the window entirely overlapping the variable V1-V2 loop in the envelope gene was not included due to the richness of insertions and deletions, which leads to poor alignment). To increase phylogenetic resolution and accuracy, we used the phyloscanner options to merge overlapping paired-end reads into single, longer reads, and to delete drug resistance sites (Gatanaga et al. 2002; Johnson et al. 2011; Wensing et al. 2015) which are known to be under convergent evolution.

Figure 4 shows the resulting phylogenies for four windows, chosen for clarity when visually inspected. The phylogenies illustrate single infection (patient A), dual infection (patient B), contamination (from the sample of patient C to the sample of patient D) and transmission (from patient E to patient F, possibly via an unsampled intermediate individual). Colouring on each phylogeny illustrates host subgraphs.

**Figure 4:**
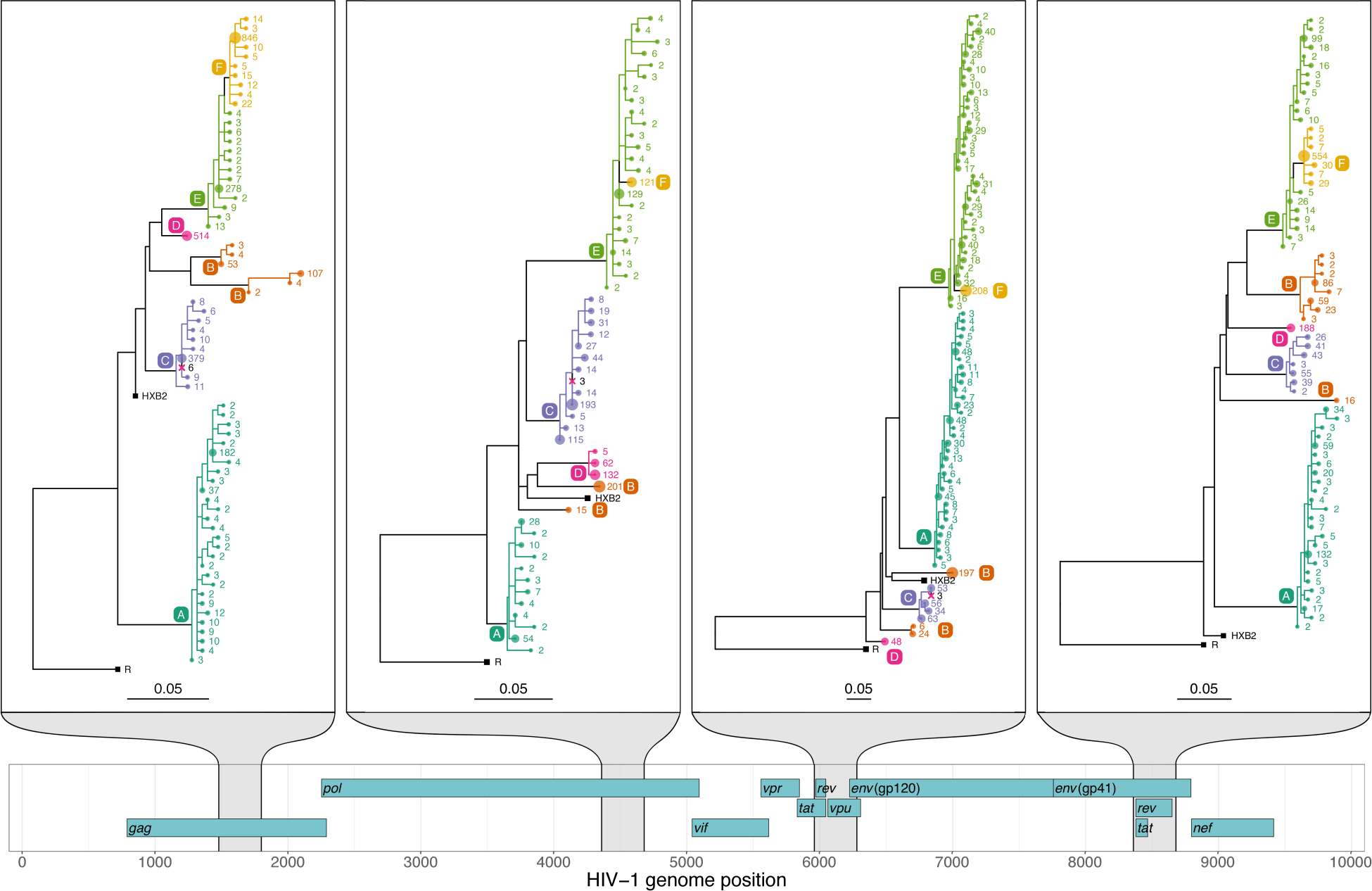
[phyloscanner analysis of four illustrative windows of the HIV-1 genome. A map of the HIV-1 genome is shown at the bottom with the nine genes in the three reading frames. Phylogenies are shown for the four windows highlighted in grey, with scale bars measured in substitutions per site. Tip labels are coloured by patient, as are all nodes assigned to that patient by ancestral reconstruction, and the branches connecting these tips and nodes; a solid block of colour therefore defines a single subgraph for one patient (see main text). The number labelling each tip is the number of times that read was found in the sample, and the size of the circle at each tip is proportional to this count. The count is after merging all identical reads and reads differing by a single base pair (merging similar reads can be done for computational efficiency, or as here, for presentational clarity). External references included for comparison are shown with black squares. One is HXB2; the other, labelled R, is a subtype C reference used to root each phylogeny. The six patients are labelled A through F. **Single infection:** patient A is a singly infected; all reads from this patient form a single subgraph. **Dual infection:** patient B is inferred to be dually infected, as is apparent by the fact that ancestral reconstruction produces two subgraphs in each window. **Contamination:** patients C and D are both singly infected, but we infer that some contamination has occurred from C to D. Patient D's sample has a small number of reads that are identical to reads from patient C, but much less numerous. Such reads are removed, but are shown here as crosses in the clade of patient C, for illustrative purposes. **Transmission:** in all four windows shown here, the reads of patient F are seen to be wholly descended from within the subgraph of reads of patient E. We infer that patient E infected patient F, either directly, or indirectly via an unsampled intermediate. Patient F having a single subgraph that is linked to patient E by a single branch indicates that the viral population was bottlenecked down to a single sampled ancestor during transmission.]

#### Contamination

Filtering for contamination is an important part of analysis of NGS data. Contamination may be physical contamination of one sample into another, or low-level barcode switching which occurs during the multiplexing and demultiplexing steps which are central to the high throughput of NGS. phyloscanner uses two criteria to identify reads as likely contaminants (either criterion is sufficient). The first is that they are exact duplicates of reads from another patient, but much less numerous; the second is that they form an additional host subgraph separated from the primary subgraph, but with too few reads to a call of multiple infection. This The second means that the source of the contaminant reads need not be present in the analysed dataset to infer contamination. These reads are flagged according to tuneable parameters (which will depend on the precise sample and method used), and blacklisted from further analysis (marked by pink crosses in Figure 4). We note that in general, phylogenetic patterns associated with transmission are distinct from those associated with contamination: the process of transmission is accompanied by within-host evolution in the recipient, whereas contamination is not.

#### Multiple infections

If the phylogeny and host-state reconstruction are correct, the number of subgraphs a patient has equals the number of founder pathogen particles with sampled descendants (for example if this is 2, a dual infection is inferred). Sampling effects mean that representatives of these multiple infections may not be present in all windows.

#### Transmission

Nodes of the phylogeny not in any patient's subgraph are coloured black in our figures, as are branches connecting nodes not part of the same subgraph. These black regions connect the different host subgraphs to each other, and so correspond to the pathogen jumping between hosts; each region must contain one or more transmission events. They may, or may not, correspond to the passage of the pathogen lineage through one or more unsampled hosts. The probability of an indirect transmission will increase with the size of the black region and may be best investigated by examining the subgraph relationships and branch lengths together.

#### Genome-wide summary statistics

In general, a phyloscanner analysis may produce a large number of phylogenies and associated ancestral reconstructions. These can be output both as annotated NEXUS format files, and as PDF files created with ggtree (Yu et al. 2017) for rapid visual inspection. Statistics are calculated to summarise the wealth of information in the phylogenies; these are shown for the 6 patients and 54 genomic windows in Figure 5. They include measures of within-host diversity, measures that allow rapid identification of multiply infected individuals, and a basic metric of recombination (defined in the supplementary section S3).

**Figure 5.**
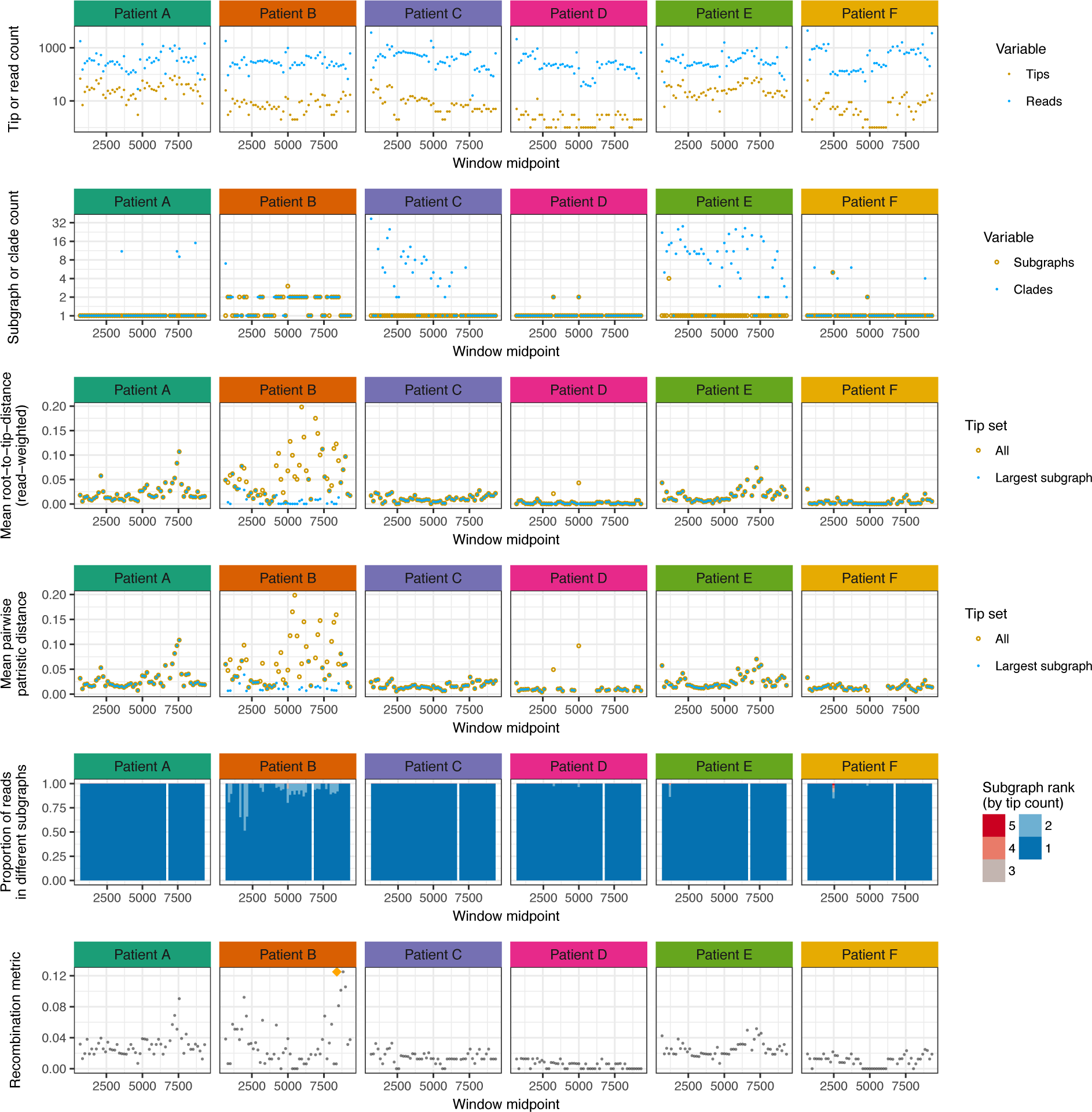
Summary statistics for six illustrative HIV-1 infected patients. Each column shows data from a single patient; each row is one or two statistics, plotted along the genome. **Top row:** number of reads, and number of unique reads (corresponding to tips in the phylogeny). **Second row:** the number of clades required to encompass all and only the reads from that patient, and the number of subgraphs (see Fig. 3 for clarification of these quantities). In many windows, though not all, the reads of patient B form two subgraphs: evidence of dual infection. For patients C and E, we see a single subgraph but many clades. This is because of the presence of reads from other patients (D and F, respectively, as seen in Fig. 4) inside what would otherwise be a single clade, turning a monophyletic group into polyphyletic group (which requires splitting in order to form clades). **Third row:** within-host divergence, quantified by mean root-to-tip distance. Defining a patient's subtree as the tree obtained by removing all tips not from this patient, we calculate root-to-tip distances both in the whole subtree and in just the largest subgraph. For patient B, this distinction is substantial due to the very large distance (~0.1 substitutions/site) between the two subgraphs of this dually infected patient. For singly infected patients, divergence may correlate with time since infection. **Fourth row:** for each window, a stacked histogram of the proportion of reads in each subgraph. For patient B, when two subgraphs are present, an appreciable proportion of reads are in the second one (mean 12%). The histogram is absent in the window that was excluded by choice. **Bottom row:** a score based on Hamming distance (between 0 and 1) of the extent of recombination in that window. The highest score across all six patients and all windows is indicated with an orange diamond; the reads giving rise to this score are shown in supplementary Figure S6.]

In a single window, phyloscanner classifies two patients to be related if they are adjacent (see supplementary section SI) and optionally, also “close”, i.e. that their subgraphs are within a prespecified patristic distance of each other. Relationships are further categorised by the ancestry, or lack of it, that is suggested by the tree topology. To summarise transmission across all windows, phyloscanner output summarises the number of windows in which each pair of patients are related, and the topological nature of that relationship. This allows the complete set of relationships between all patients in the dataset to be visualised in graph form. For example, in this dataset, only two of the six patients, E and F, are related in at least half of the windows. In Figure 6A the counts of the different topological relationships between these two patients are displayed. With many links between many patients these graphs become difficult to interpret visually; a threshold on the number of windows for links to be displayed is therefore helpful. phyloscanner also produces a second version of the graph simplified further, shown in Figure 6B. Here a single link appears if relatedness of any type is present in 50% of windows, and that link is an arrow if transmission in that direction is inferred in at least 33% of windows. (The 50% and 33% thresholds are defaults that can be changed.) These relationship diagrams were plotted using Cytoscape 3.5.1 (Shannon et al. 2003).

**Figure 6.**
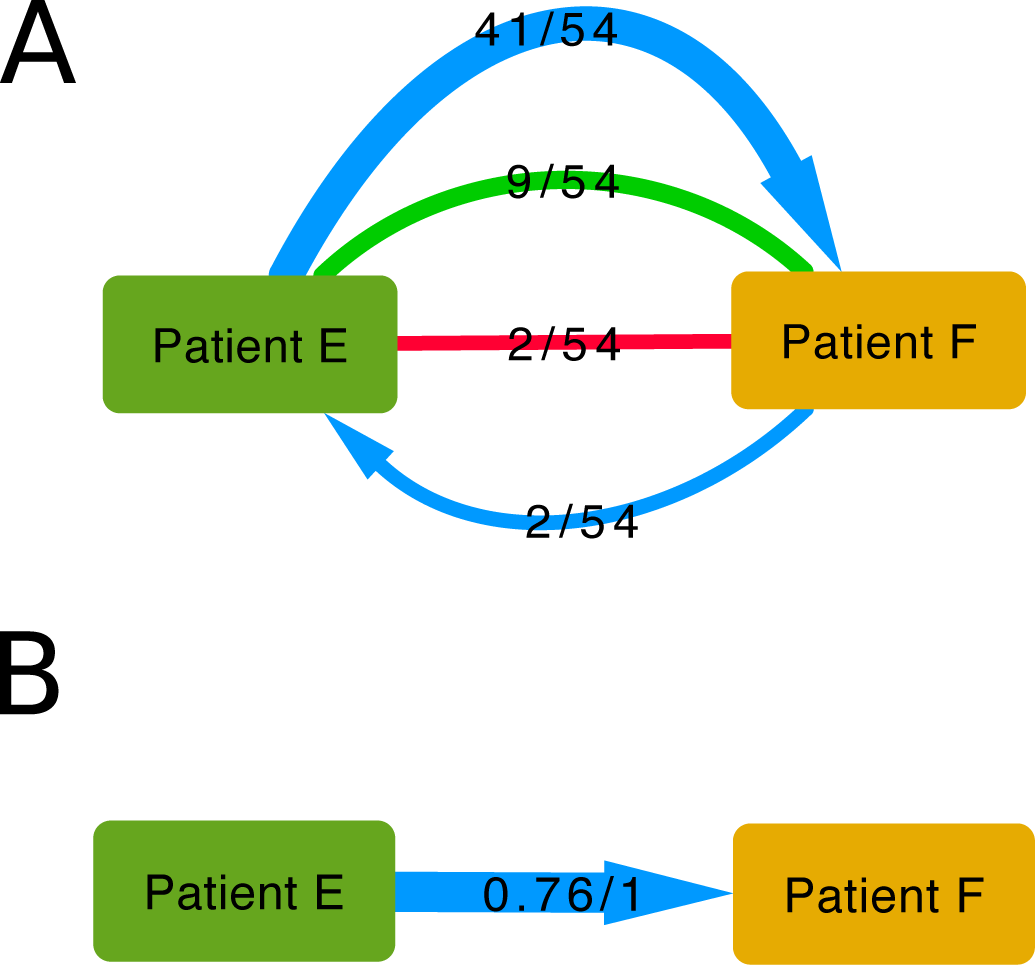
[– Relationship graphs: visual representations of the relationship between two connected patients infected with HIV-1. The power of phyloscanner in studying transmission events comes from aggregating information over many within- and between-host phylogenies, in this case obtained from different windows of the whole HIV-1 genome. In the top diagram, the outcomes from all 54 windows are shown. The top blue arrow shows that in 41 windows, patient E was inferred to be ancestral to patient F, with a single bottleneck. The bottom blue arrow shows that in 2 windows the reverse was true – F was ancestral to E. The undirected red line shows that in 2 windows, the patients were linked by “complex” ancestry, with the direction unclear. The undirected green line shows that in 9 windows the patient subgraphs were adjacent and close, but no ancestry was implied by the topology. In no window was transmission of more than one lineage inferred, and in no window were the patients distant and unlinked. (See supplementary section SI 1 for more details on these categories.) A simplification of these relational data is shown in the bottom diagram, with a single directed arrow. The first number indicates the proportion of windows supporting transmission in the direction of the arrow, and the second number indicates the proportion of windows supporting transmission in either direction.]

Diagrams such as those in Figure 6, when extended to greater numbers patients, will not always represent a single, coherent transmission tree amongst all the patients in the dataset (as can be seen in Figures 7 and 9). Instead, they simply summarise each pairwise relationship. As a result, we refer to them as “relationship graphs”. The inference of a single, most probable transmission tree over all windows is complicated by the presence of multiple infections, incomplete transmission bottlenecks, and missing data for some patients in some windows. To our knowledge, no method yet exists to produce a consensus transmission history that takes into account all these possibilities.

**Figure 7.**
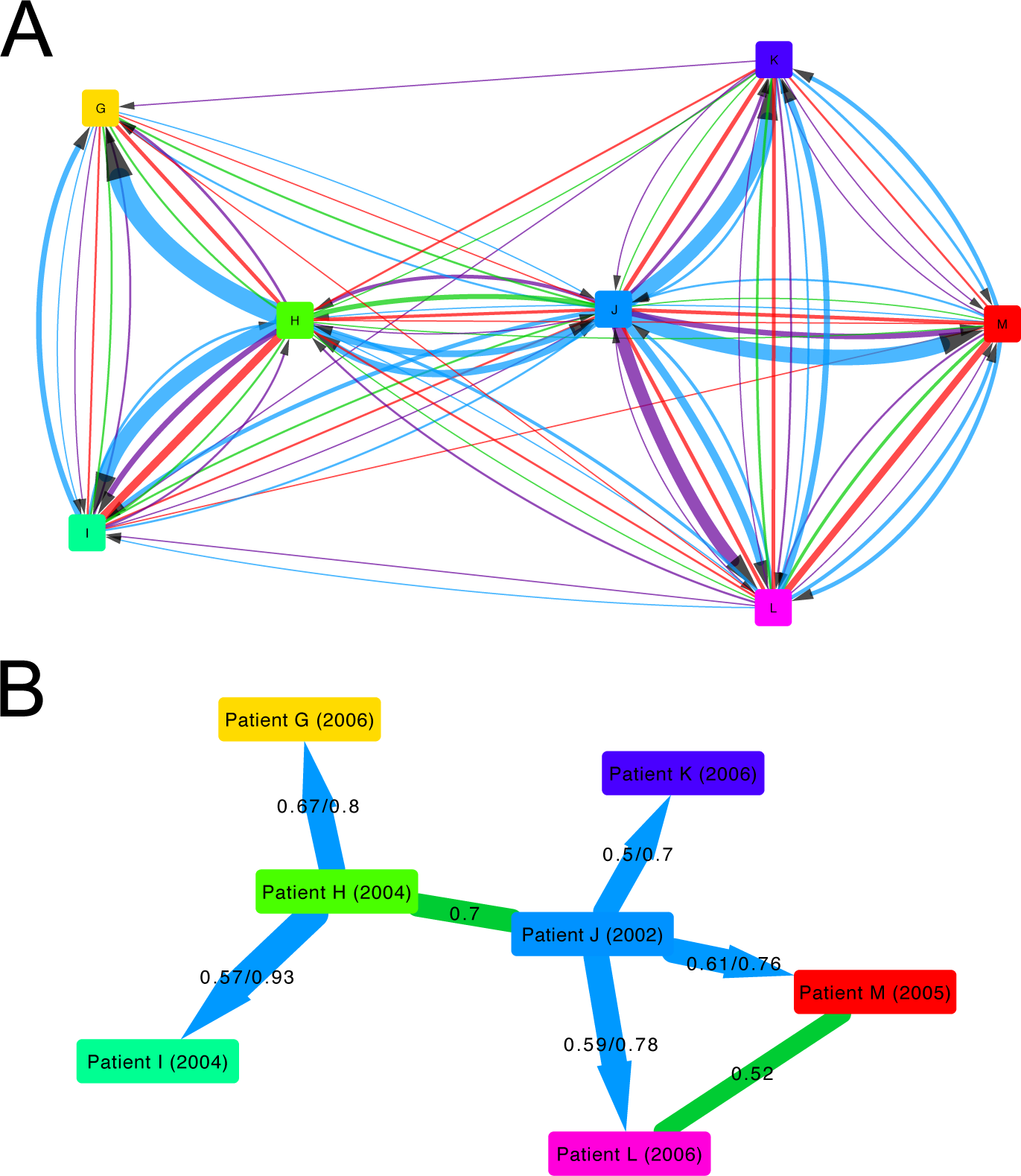
[– The relationship between 7 patients infected with HIV-1. The colouring and numbers on the arrows connecting patients are as in Figure 6; in addition, the lower diagram here contains undirected green lines as well directed blue lines. These green lines suggest that the pair are close in the transmission network but with unknown transmission direction; the single number on the line indicates the proportion of windows supporting this. The known or estimated year of infection is shown in parentheses after each patient's label.]

### Resolving the transmission pathway within a HIV-1 phylogenetic cluster

To illustrate the resolution into the transmission process that can be obtained by phyloscanner, we chose a set of 7 patients from the BEEHIVE study that were found to be closely connected in the chain of transmission (Fig. 7). 3 of the patients' samples were sequenced with Illumina MiSeq and 4 with Illumina HiSeq; the resulting reads were processed and mapped using IVA and shiver as previously, with the mapped reads given as input to phyloscanner. phyloscanner summarises all the pairwise relationships between individuals in each window (Figure 7A), suggesting a complex network. However, we find that when we focus on the most likely inferences of source attribution (Figure 7B), phyloscanner largely resolves a complex set of pairwise relationships into a coherent transmission network, that is consistent with the years of seroconversion. However, this is not guaranteed to be the case: an exception is the triangle connecting Patients J, L and M, where there is too much uncertainty in the relationships amongst the triplet to resolve their ancestry.

### HIV-1 sequenced with Roche 454

A subset of patients from the BEEHIVE study were also sequenced using the Roche 454 platform; results from their analysis with phyloscanner are in Supplementary Information section SI 2.

### HCV sequenced with Oxford Nanopore MinION

To further illustrate phyloscanner's applicability to different sequencing platforms and also different pathogens, we used it to analyse HCV viral data sequenced using the Oxford Nanopore MinION device. Plasma samples were obtained from four patients in the BOSON study (Foster et al. 2015), a phase 3 randomized trial of antiviral therapy with sofosbuvir (trial registration NCT01962441). Sequencing was performed using RNAseq-based methods previously described for Illumina (Bonsall et al. 2015) and adapted for the MinION device. Briefly, plasma-derived RNA was reverse transcribed, then sequencing libraries were prepared for each sample using Oxford Nanopore adapters and customised barcoded primers. These were pooled and enriched using HCV-specific nucleotide baits before sequencing on a MinION R9.0 flow cell. Viral sequences were identified and mapped using BLASTN (Altschul et al. 1990), standard reference sequences and BWA (Li and Durbin 2009). See Materials and Methods for more details. The resulting BAM files were used as input for phyloscanner, with a window size of 600 bp and no overlap between windows. Nanopore sequencing platforms are capable of producing longer inserts than those of Illumina, at the cost of a higher error rate (approximately 10% erroneous base calls). Despite this error, phyloscanner could phylogenetically resolve the within- and between-host evolution, shown in Figure 8.

**Figure 8.**
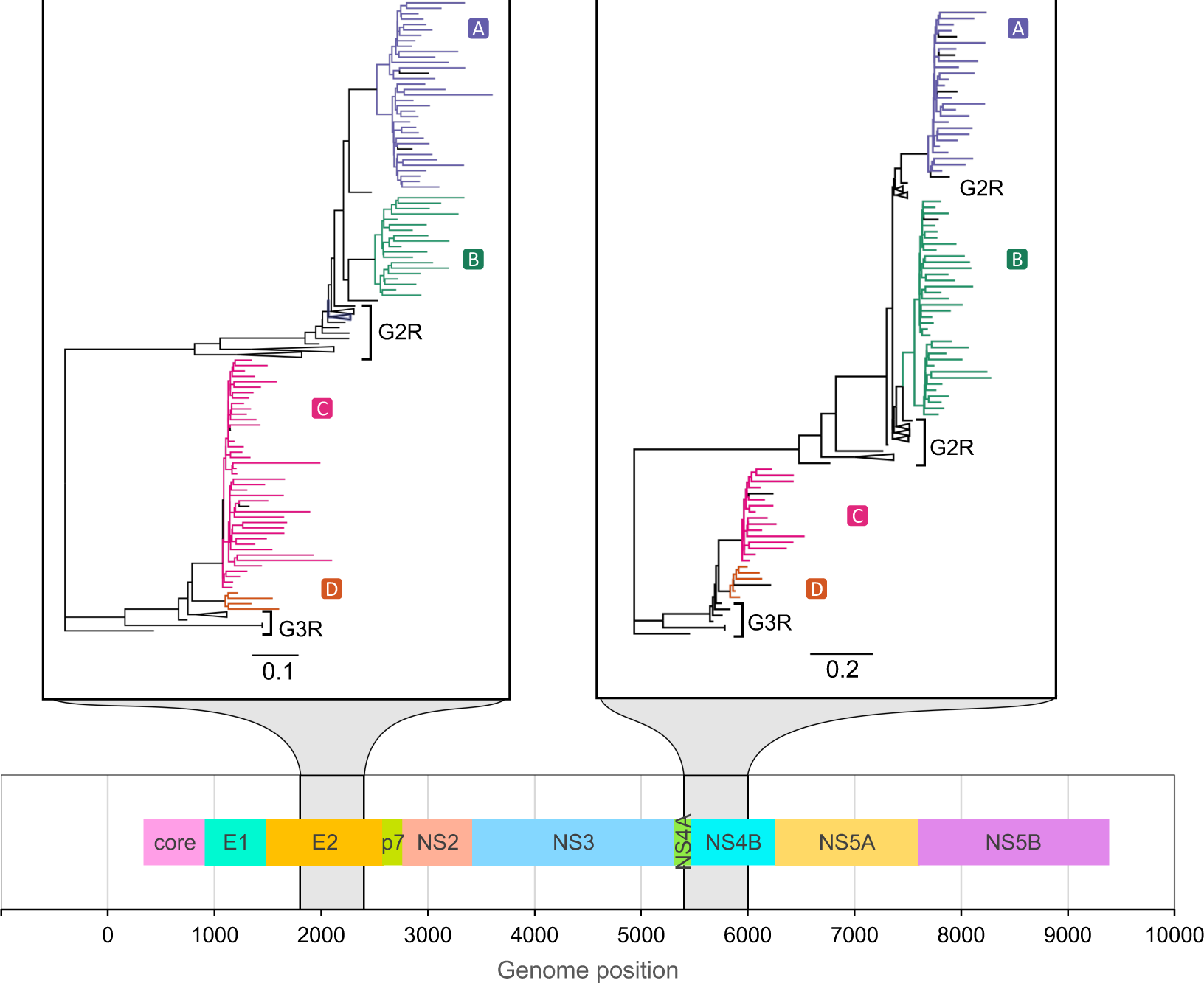
[phyloscanner analysis of two illustrative windows of the HCV genome. Sequence data from four individuals was obtained with the Oxford Nanopore MinION device. A continuous region of the phylogeny with the same colour shows a subgraph for one patient (see main text). Black tips were flagged as contamination and excluded. Patient-derived sequences clustered with respective genotype 2 and genotype 3 references (G2R, G3R) as expected from the virus genotypes known from the clinical information available for participants. Two windows, 600 bp in length, are shown for the E2 and NS4B genes at positions given by the genome map (bottom panel).]

### Multiple colony picks per carrier of *S. pneumoniae*

phyloscanner's analysis of phylogenies need not be restricted to those derived from deep sequencing data in different windows of the genome: it can also be applied to datasets where within-host diversity is captured by SGA or sequences from multiple colony picks per individual. We illustrate this approach with the *S. pneumoniae* data of Croucher et al. (Croucher et al. 2016), specifically the BC1-19F cluster. This dataset consists of 286 sequences from 92 individuals carrying the bacterium (with multiple colonies per carrier). These were sequenced with Illumina HiSeq, though for SGA data sequencing platform is largely irrelevant to interpretation, since each sequenced sample should not contain any real within-sample diversity by design. Genomes were processed with Gubbins (Croucher et al. 2015) to remove substitutions likely to have been introduced by recombination. As each of these sequences is a whole genome (unlike the short reads produced by NGS), we did not split the genome into windows to be analysed separately. Instead, we represented phylogenetic uncertainty by generating a posterior set of 100 phylogenies using MrBayes 3.2.6 (Ronquist et al. 2012) and analysed these with phyloscanner. Ancestral state reconstruction was performed on each posterior phylogeny independently, relationships between carriers identified, and the results summarised over the entire set. In each phylogeny, carriers were inferred as being related if the minimum patristic distance between two nodes from the subgraphs associated with each was less than 7 substitutions and they were categorised as adjacent (explained in Supplementary Information section SI 1.5). This distance threshold was selected to demonstrate the method as it picked out obvious clades in the phylogeny as groups, and was not chosen to imply direct transmission. Retaining such relationships where they existed in at least 50% of posterior phylogenies revealed 18 separate groups of carriers whose bacterial strains were closely related (see Fig. 9).

**Figure 9.**
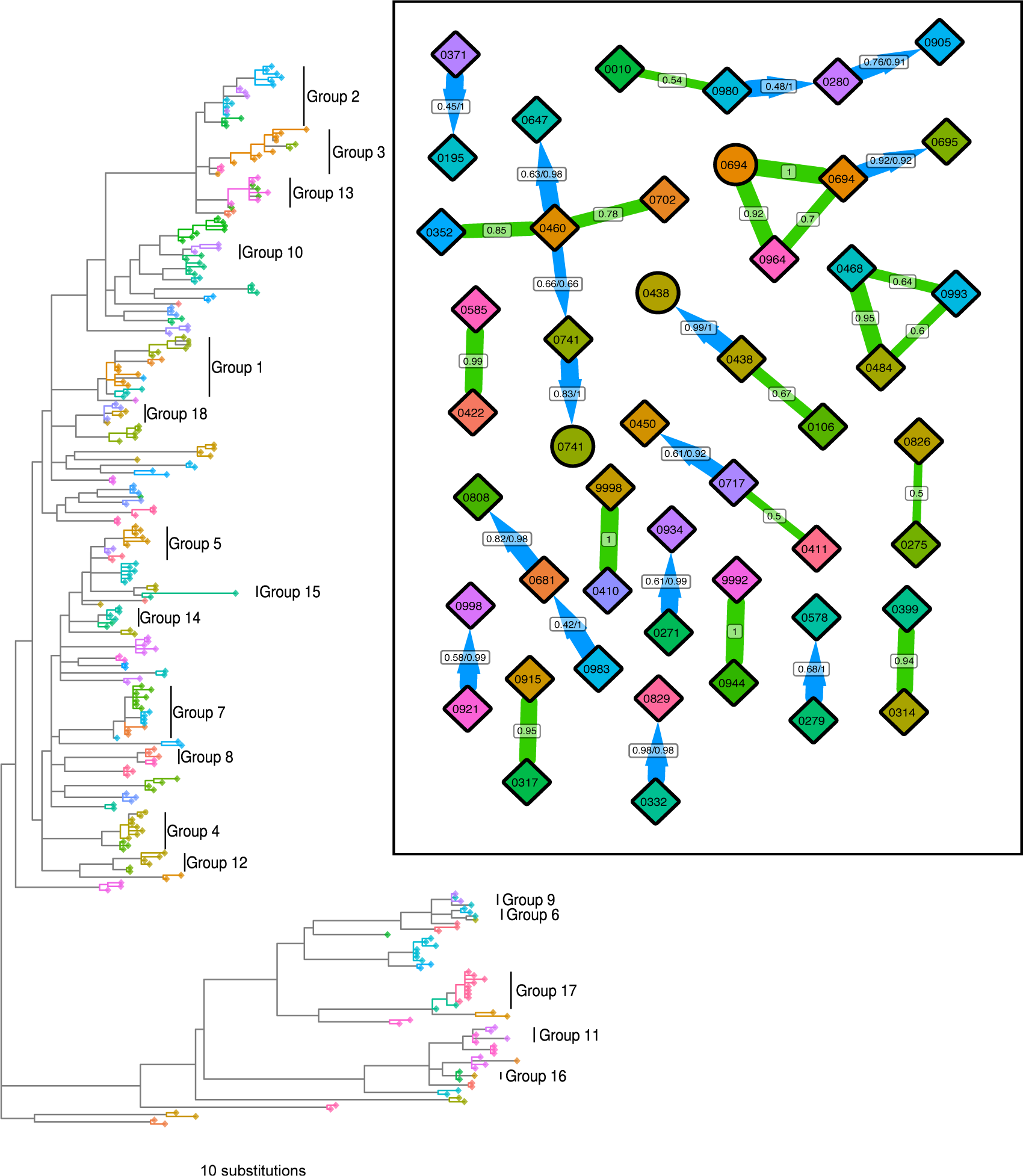
[– Phylogeny and relationships between *S. pneumoniae* carriers. The phylogeny shown is the MrBayes consensus tree. Tip shapes are coloured by carrier, with mother and infant pairs sharing the same colour; diamonds represent infants and circles mothers. All nodes assigned to a carrier by ancestral reconstruction, and the branches connecting these tips and nodes, are given the same colour as that carrier's tips; a solid block of colour therefore defines a single subgraph for one carrier (see main text). Regions of the phylogeny not in any carrier's subgraph are grey. These regions connect carriers' subgraphs to each other, and so each must contain one or more transmission events. The carrier relationship diagram (inset) displays the relationships between the carriers in 18 identified groups, in the same fashion as in Figures 6 and 7, except that here the numbers represent the proportion of phylogenies from the posterior set, rather than the proportion of genomic windows in which both patients have sequence data. The clades representing these 18 groups are labelled in the phylogeny.]

Note that if some residual signals of recombination remain after processing with Gubbins, analysing the full-length genomes in windows by choice (rather than by necessity, as with short-read NGS data) could mitigate this effect at the cost of reduced phylogenetic resolution in each window. The merits of this could be explored in a dedicated analysis of such a dataset; here we simply illustrate application of phyloscanner to full-length sequences as opposed to genomic windows.

## Discussion

Improving our understanding of the transmission of pathogens is valuable for identifying epidemiological risk factors – the first step for targeting public health interventions for efficient impact. Phylogenetic analysis of one pathogen sequence per infected individual may identify clusters of similar sequences that are expected to be close in a transmission network. However, nothing is learned about the direction of transmission within the network. Indeed it may be that none of the individuals transmitted the pathogen to anyone else, and they were all infected by a common individual who was not sampled. Through automatic fitting of maximum-likelihood evolutionary models to within- and between-host genetic sequence data, phyloscanner enhances resolution into the pathogen transmission process. An evidence base is built up by analysing many phylogenies, notably through consideration of NGS reads in windows along the pathogen's genome. The relationship between infected individuals is no longer quantified by a single number summarising closeness, but by a rich set of data resulting from ancestral host-state reconstruction for each phylogeny.

Romero-Severson *et al.* (Romero-Severson et al. 2016) demonstrated the utility of parsimony for the assignment of ancestral hosts to internal nodes in a phylogeny containing many tips from two infected individuals, for simulated HIV-1 data. We have continued with this approach, developing it for suitability for real sequence data from many infected individuals. In particular we allow for (i) contamination, (ii) multiple infections, and (iii) the possible presence of unsampled hosts in the tree. Details of two such parsimony algorithms, available for use in phyloscanner, are presented in the supplementary section SI 1. Parsimony has the advantage that a reconstruction can be completed in reasonable computational time even for phylogenies with tens of thousands of tips. Other methods of reconstructing the host state of internal nodes could also be suitable and may be added to the package in future. Our identification of contamination and multiple infections is highly valuable in its own right: the former because this is critical for any empirical study of within-host diversity, and the latter because such individuals may be special cases clinically and for pathogen evolution. Transmission of multiple distinct pathogen strains may occur simultaneously, or sequentially – 'super-infection’. phyloscanner can detect both cases, though distinguishing them is difficult without longitudinal sampling (it could be possible through inference of timed trees, or using the diversity of each separate infection as a proxy for its age).

Great care must be taken to correctly interpret the ancestry of pathogens infecting individuals. Even if ancestry were established beyond any doubt, individual X's pathogen being ancestral to individual Y's pathogen does not imply that X infected Y: the pathogen could have passed through unsampled intermediate hosts. Nevertheless the ancestry does provide valuable epidemiological information, as X has been identified as a transmitter (and Y a recipient not far down the same transmission chain). Finding likely transmitters in a large population cohort would allow risk factors to be identified and quantified.

Furthermore, inference of ancestry is itself subject to uncertainty. The inference of ancestry depends on the correct rooting of the phylogeny, in order that the direction in which evolution proceeded over time is known. Molecular clock analyses (such as implemented in TempEst (Rambaut et al. 2016)) can aid correct rooting when the sampling dates of the tips of the phylogeny are known.

The relationships between infected individuals are inferred by phyloscanner across many phylogenies, for example those constructed from NGS reads in windows along the pathogen genome. By analysing many phylogenies, phyloscanner mitigates the effect of random error - any error that is independent in each phylogeny. We therefore give greater credibility to those relationships observed many times than to those observed only once. However, systematic error may arise, for example, due to different patients being sampled at different stages of infection, with different amounts of within-host diversity to analyse (Romero-Severson et al. 2016). Given uncertainties in any individual assignment, we recommend phyloscanner for population-level analyses, rather than focussing on isolated transmission events (as we have done here, for simplicity in explaining the method).

The fraction of genomic windows in which a given relationship is inferred between individuals (for example A infecting B directly or indirectly), is not equal to the probability of that relationship being true. However it provides a measure of the robustness with which the available data support that conclusion. This is analogous to bootstrapping – sampling with replacement from the same sequence alignment, to create a set of similar phylogenies. Here however, different windows of the genome make use of different sequence data. Given the potential for disagreement between different windows due to genuine biological variation, imperfect sequencing procedures etc., agreement between a fraction *x* of (non-overlapping) windows is a stronger statement of robustness than agreement between a fraction *x* of bootstraps. Identification of transmission events with phyloscanner will involve false positives and false negatives; these will be context dependent, depending on how strictly transmission thresholds are defined (which balance sensitivity and specificity) and on the inclusion of sequences similar to those being investigated. We will illustrate this in two works in preparation examining large population studies.

Whilst our emphasis has been on extracting broad-brush information from the rich within-and-between host phylogenies, these phylogenies contain more information that could be used in future research. A specific example is that by resolving the transmission event at a finer level of genetic detail, it is possible to identify which pathogen genotypes are typically transmitted and which ones are not, with potential relevance for vaccine design.

By providing a tool for automatic phylogenetic analysis of NGS deep sequencing data, or multiple genotypes per host generated by other means, we aim to simplify identification of transmission, multiple infection, recombination and contamination across pathogen genomics.

## Materials and Methods

### Generation and assembly of the BEEHIVE Illumina data

Viral RNA was extracted manually from blood samples following the procedure of Cornelissen *et al.* (Cornelissen et al. 2016). RNA was amplified and sequenced according to the protocol of Gall *et al.* (Gall et al. 2012; Gall et al. 2014). Briefly, universal HIV-1 primers define four amplicons spanning the whole genome. 5 μl of amplicon I was pooled with 10 μl each of amplicons II–IV. Libraries were prepared from 50 to 1000 ng DNA as described in Quail et al. (Quail et al. 2008; Quail et al.), using one of 192 multiplex adaptors for each sample. Paired-end sequencing was performed using an Illumina MiSeq instrument with read lengths of length 250 or 300 bp, or in the ‘rapid run mode’ on both lanes of a HiSeq 2500 instrument with a read length of 250 bp.

For each sample, the reads were assembled into contigs using the *de novo* assembler IVA. The reads and contigs were processed using shiver as described previously (Wymant et al. 2016). In summary: non-HIV contigs were removed based on a BLASTN search against a set of standard whole-genome references (Kuiken et al. 2012). Remaining contigs were corrected for assembly error then aligned to the standard reference set using MAFFT (Katoh et al. 2002). A tailored reference for mapping was then constructed for each sample using the contigs, with gaps between contigs filled by the corresponding part of the closest standard reference. The reads were trimmed for adapters, PCR primers and low-quality bases using Trimmomatic (Bolger et al. 2014) and fastaq (https://github.com/sanger-pathogens/Fastaq). Contaminant reads were removed based on a BLASTN search against the non-HIV contigs and the tailored reference. The remaining reads were then mapped to the tailored reference using SMALT (<u>http://www.sanger.ac.uk/science/tools/smalt-0)</u>.

### Generation and assembly of the HCV Oxford Nanopore MinION data

Viral RNA was extracted from plasma using the NucliSENS^®^ easyMAG^®^ total nucleic acid extraction system (Biomerieux) and sequencing libraries were prepared using a modified version of an RNA-seq based protocol with a virus enrichment step. Briefly, the NEBNext^®^ Ultra™ Directional RNA Library Kit (New England Biolabs, Ipswich, MA, USA) was used to generate cDNA from 5ul of total RNA. The NEBNext^®^ Ultra™ II End Repair/dA-Tailing Module and Blunt/TA Ligase (New England Biolabs, Ipswich, MA, USA) were used for end repair of dsDNA and ligation of PCR adapters (Oxford Nanopore Technologies) to allow for 18 cycles of PCR using custom barcoded primers with a post-PCR clean-up with 1x Ampure XP (Beckman Coulter, Pasadena, CA, USA). Each library was quantified by Quant-iT™ Qubit^®^ dsDNA HS Assay Kit and size distribution analysed using Agilent Tapestation High Sensitivity D5000 ScreenTape System. Approximately equimolar quantities of each library were pooled to a total of 500 ng mass and processed for probe enrichment using customized xGen^®^ Lockdown^®^ 120mer probes specific to HCV (Integrated DNA Technologies, Inc., Coralville, Iowa, USA) and a modified Roche NimbleGen protocol for hybridization of amplified sample libraries with a shorter 4 hours hybridization time and on-bead post-enrichment PCR (12 cycles). The enriched pool was prepared for sequencing on a MinION R9.0 flow cell using the SQK-NSK007 2d ligation kit. Raw fasta5 sequence files were base called and demultiplexed using Metrichor software. Viral sequences were identified and trimmed using a BLASTN search of the Los Alamos database of HCV genotype references (Kuiken et al. 2005), then mapped to the closest matching reference using BWA (with the command bwa mem −x ont2d). Consensus sequences were called from the bam files and used as references for a second iteration of read mapping.

### The phyloscanner Method

For application of phyloscanner to deep sequence NGS data, the required input is a set of files in BAM format (Li et al. 2009) each containing the reads from one sample that have been mapped to a reference, and a choice of genomic windows to examine. A sensible choice of windows would normally tile the whole genome, perhaps skipping regions that are rich in insertions and deletions (leading to poor sequence alignment). Windows should be wide enough to capture appreciable within-host diversity, but short enough for some reads to fully span them; options in the code help to inform the user's choice. There is no lower limit to the length of reads given as input, however as read length decreases, phylogenetic resolution will suffer. phyloscanner determines the correspondence between windows in different BAM files by aligning the mapping references in the BAM files. Using the same reference for mapping all samples would negate the need for this step, but it is of paramount importance to tailor the reference to each sample before mapping to minimise biased loss of information (Wymant et al. 2016). For each window in each BAM file, all reads (or inserts, if reads are paired and overlapping) fully spanning the window are extracted using pysam (<u>https://github.com/pysam-developers/pysam</u>) and trimmed to the window edges, then identical reads are collapsed to a single read, giving a set of unique reads each with an associated count (i.e. the number of reads with identical sequence). A basic metric of recombination is calculated by maximising, over all possible sets of three sequences and all possible recombination crossover points, the extent to which one of the three sequences resembles one of the other two sequences more closely on the left and resembles the other sequence more closely on the right. Further detail is provided in the supplementary section SI 3. In each window, each sample's set of unique reads is checked against every other sample's set, with exact matches flagged to warn of between-sample contamination in the analysed dataset; all unique reads are then aligned with MAFFT, and a phylogeny is inferred with RAxML (Stamatakis 2014).

phyloscanner contains many options to customise processing and maximise the information extracted from reads and phylogenies. Standard reference genomes can be included with the reads for comparison. User-specified sites can be excised to mitigate the effect of known sites under selection on phylogenetic inference. Greater faith can be placed in the reads by trimming low-quality ends and wholly discarding reads that are low-quality, improperly paired, or rare. Reads in the same sample that differ from each other by less than a specified threshold can be merged into a single read to increase the speed of downstream processing. Overlapping paired reads can be merged into a single longer read for greater phylogenetic resolution. Every option of RAxML can be passed as an option to phyloscanner, for example specifying the evolutionary model to be fitted, or multithreading.

Optionally, the user may skip inference of phylogenies from files of mapped reads, and instead directly provide as input a phylogeny or a set of phylogenies generated by any other method.

To analyse phylogenies, phyloscanner required that they are rooted. This can be done manually, or if the phylogenies were constructed by phyloscanner from mapped reads, rooting can be achieved by providing one or more additional reference sequences with the mapped reads, and choosing one of these to use as an outgroup. The outgroup should be sufficiently distant from all sampled isolates that we can assume the most recent common ancestor of it and every isolate (i.e. the root of the whole tree) was not present in any of the sampled individuals.

Each phylogeny analysed is annotated with a reconstruction of the transition process using a modified maximum-parsimony approach to assign internal nodes to hosts or to an extra “unassigned” state. The latter is given to lineages that either must have infected a host outside the dataset, or to those where the situation is sufficiently ambiguous that this cannot be ruled out. An important parameter of the reconstruction, designated *k*, is used to help identify dual infections and contaminants. It acts as a penalty, in the parsimony algorithm, for the reconstruction of single infections showing unrealistic within-host diversity. A suitable value of *k* will depend on the pathogen under study, but as a rule of thumb, we suggest estimating a level of pairwise genetic diversity that it would be unrealistic to see in an infection from a single source, and using the reciprocal of this for *k*. In situations where the phyloscanner user is confident that dual infections and contaminants are not present, *k* can be set to zero, in which case no penalty for within-host diversity is applied.

The results of the reconstruction can be represented as a visualisation of the partial pathogen transmission tree by the process of ‘collapsing’ each subgraph (i.e. each set of adjacent nodes with the same reconstructed host; see supplementary Fig. S3) into a single node of a new tree structure. This “collapsed tree” is then analysed to identify relationships between each pair of infected individuals, according to the following categories:

- 1. Minimum distance: what is the smallest patristic distance between a phylogeny node assigned to one host and a node assigned to the other?
- 2. Adjacency: is there a path on the phylogeny that connects the two individuals' subgraphs without passing through a third individual? (“Unassigned” nodes do not interrupt adjacency.)
- 3. Topology: how are the regions from each individual arranged with respect to each other? (See supplementary Fig. S4.)

Combinations of these properties can be used to develop criteria which identify individuals who are closely linked in the transmission chain. For example, two individuals that are adjacent and within a suitable distance threshold are likely to be either a transmission pair, or infected via a small number of unsampled intermediaries. If the distance between subgraphs is large, on the other hand, separation by unsampled hosts in the chain of transmission is likely even if they are adjacent. The nature of the topological relationship between them may suggest a direction of transmission, or be equivocal.

An individual having multiple subgraphs suggests multiple infection, with the ancestor node of each subgraph inferred to be a distinct founder pathogen particle (the ancestor of that sampled subpopulation). It can be difficult to distinguish a dual infection from a sample that has been contaminated by another sample not present in the current data set (i.e. where contamination is not visible as exact duplication of another individual's read). For NGS data we make the distinction in each phylogeny based on thresholds on read counts: outside of the subgraph containing the greatest number of reads, any additional ('minor’) subgraph is designated as contamination and ignored if the number of reads it contains is below an absolute threshold, or below a threshold relative to the read count in the largest subgraph. By default, minor subgraphs with read counts exceeding both thresholds are kept, providing evidence for the presence of multiple distinct subpopulations in that genomic window. (Alternatively, a phyloscanner option allows all minor subgraphs to be entirely removed from consideration). Zanini et al. (Zanini et al. 2015) discarded reads suspected of being contamination by calculating each read's Hamming distance from the consensus, plotting the distribution of these distances, and discarding reads giving rise either to a second peak or to a ‘fat tail’ (taken to be recombinant reads). This approach is not appropriate when the data set may contain multiply infected individuals, for example for a dual infection we wish to keep the reads from each of two distinct groups that may be separated by a large distance.

### The phyloscanner Code

phyloscanner is freely available at <u>https://github.com/BDI-pathogens/phyloscanner</u>. It is written in Python and R, but can be run from the command line so that no knowledge of either language is required. Inference of within- and between-host phylogenies from BAM-format mapped reads is achieved with a single command of the form phyloscanner_make_trees.py ListOfBamsAndRefs.csv --windows 1,300,301,600,… where ListOfBamsAndRefs.csv lists the BAM files to be analysed and the fasta-format references to which the reads were mapped, and the --windows flag above specifies analysis of the genomic windows with coordinates 1-300, 301-600,…Analysis of those trees is achieved with a single command of the form phyloscanner_analyse_trees.R TreeFiles OutputLabel [choice of ancestral state reconstruction].

Included with the code is simple simulated HIV-1 data for ease of immediate exploration of phyloscanner. Within-host evolution was simulated using SeqGen (Rambaut and Grassly 1997); each resulting sequence was then converted into error-free fragments that were mapped back to the founding sequence, giving BAM-format files suitable as input for phyloscanner. We also created BAM-format files by using shiver to process publicly available HIV-1 reads sequenced with Illumina MiSeq. A tutorial walking the user through a simple application of phyloscanner to the simulated data, and a more sophisticated application to this real public data, is available from the GitHub repository with the code itself.

Running phyloscanner on the six HIV-1 samples presented in the first results section took 18 minutes on one core of a standard laptop, 10 minutes of which was running RAxML. A number of options allow the user to speed up phyloscanner. Firstly it is ‘embarrasingly’ parallelisable, in that each window of the genome can be processed separately (e.g. the 54 windows used for the HIV data could have been processed via 54 jobs run in parallel). Secondly all options of RAxML can be passed as options to phyloscanner, including multithreading. Thirdly the number of unique sequences kept for phylogenetic inference can be controlled through various options, notably merging of similar reads and/or a minimum read count. Fourthly the user can easily use a different tool for phylogenetic inference instead of RAxML by using the --no-trees option of phyloscanner_make_trees.py, and running the desired tool on the fasta file of processed reads that is output for each window. (As an example running FastTree(Price et al. 2009) on the same data took 28 seconds instead of the 10 minutes needed by RAxML.)

## Acknowledgments

We thank Katrina Lythgoe for helpful discussions, and Céline Christiansen-Jucht for comments on the manuscript. This work was funded by ERC Advanced Grant PBDR-339251. We acknowledge funding from Bill & Melinda Gates Foundation through PANGEA-HIV. The STOP-HCV Consortium is funded by a grant from the Medical Research Council (MR/K01532X/1). We thank Gilead Sciences for providing HCV plasma samples from the BOSON clinical study for use in these analyses. We also thank HCV Research UK (funded by the Medical Research Foundation) for their assistance in handling and coordinating the release of samples for these analyses. This work used the computing resources of the UK MEDical BIOinformatics partnership - aggregation, integration, visualisation and analysis of large, complex data (UK MED-BIO) which is supported by the Medical Research Council [grant number MR/L01632X/1].

## The BEEHIVE Collaboration

Jan Albert, Margreet Bakker, Norbert Bannert, Ben Berkhout, Daniela Bezemer, François Blanquart, Marion Cornelissen, Jacques Fellay, Katrien Fransen, Christophe Fraser, Astrid Gall, Annabelle Gourlay, M. Kate Grabowski, Barbara Gunsenheimer-Bartmeyer, Huldrych F. Günthard, Matthew Hall, Mariska Hillebregt, Paul Kellam, Pia Kivelä, Roger Kouyos, Oliver Laeyendecker, Kirsi Liitsola, Laurence Meyer, Swee Hoe Ong, Kholoud Porter, Peter Reiss, Matti Ristola, Ard van Sighem, and Chris Wymant.

Acknowledged contributors to the cohorts in the BEEHIVE Collaboration are listed in supplementary section SI 4.

## The STOP-HCV Consortium

Eleanor Barnes, Jonathan Ball, Diana Brainard, Gary Burgess, Graham Cooke, John Dillon, Graham R Foster, Charles Gore, Neil Guha, Rachel Halford, Cham Herath, Chris Holmes, Anita Howe, Emma Hudson, William Irving, Salim Khakoo, Paul Klenerman, Diana Koletzki, Natasha Martin, Benedetta Massetto, Tamyo Mbisa, John McHutchison, Jane McKeating, John McLauchlan, Alec Miners, Andrea Murray, Peter Shaw, Peter Simmonds, Chris C A Spencer, Paul Targett-Adams, Emma Thomson, Peter Vickerman, and Nicole Zitzmann.

## The Maela Pneumococcal Collaboration

Stephen D. Bentley, Claire Chewapreecha, Nicholas J. Croucher, Simon Harris, Jukka Corander, David Goldblatt, Julian Parkhill, Francois Nosten, Claudia Turner, and Paul Turner.

